# Towards an eco-epidemiological framework for managing freshwater crayfish communities confronted with crayfish plague

**DOI:** 10.1101/2024.07.31.606058

**Authors:** Claudio Bozzuto, Heike Schmidt-Posthaus, Irene Adrian-Kalchhauser, Simone Roberto Rolando Pisano

**Author notes:** Correspondence to: Simone R.R. Pisano, Institute for Fish and Wildlife Health, Vetsuisse Faculty, University of Bern, Länggassstrasse 122, 3001 Bern, Switzerland.

## Abstract

Wildlife diseases figure prominently among the main causes of biodiversity loss worldwide. Especially fungal and fungus-like pathogens are on the rise, wreaking havoc across the tree of life by threatening species persistence and destabilizing ecosystems. A worrisome example are freshwater crayfish species in Eurasia and Oceania, facing the dual challenge of introduced competitive crayfish species and an introduced water mold (*Aphanomyces astaci*) causing crayfish plague. *A. astaci* locally extinguishes susceptible native crayfish populations, while non-native individuals (mostly from North America) remain largely unaffected. Despite its significant impact and its ∼150 years of presence in Europe, studies and disease management recommendations for crayfish plague that are firmly rooted in epidemiological theory are scarce. Here, we present a practical eco-epidemiological framework to understand how multi-species crayfish communities react to crayfish plague introductions. The framework is based on the observation that the dynamics of crayfish communities are mainly determined by life-history characteristics, within- and among-species competition, effects of generalist predators (including fishing), and host-pathogen interactions. From this ecological and epidemiological context, we derive fundamental epidemiological metrics, single-host species and community-level basic reproduction numbers (*R*_0_). We investigate how host species densities affect the likelihood of a disease outbreak in a crayfish community, and we demonstrate that a community’s *R*_0_ value is simply the sum of the community’s single-host species *R*_0_ values, adjusted for competition and predation. We further demonstrate how *R*_0_ can be used to guide preventative and mitigation actions for crayfish communities. For example, we show how *R*_0_ expressions – even without a detailed parametrization – can be used to construct regional risk rankings for different crayfish communities, for an effective allocation of resources to local conservation plans. Our eco-epidemiological framework will also be of interest to the management of other aquatic host-pathogen systems with water-borne pathogen transmission as the main route of pathogen spread.

## Introduction

Emerging wildlife pathogens figure prominently among the threats to life on Earth in the Anthropocene (Fisher et al., 2012; Jaureguiberry et al., 2022; McCallum et al., 2024). For example, amphibian or plant fungal pathogens have been observed to trigger catastrophic consequences for populations and ecosystems worldwide (e.g. Karnosky, 1979; Scheele et al., 2019). Globalization has spurred pathogen emergences, since the introduction of a pathogen often happens via the transport of a carrier organism (Fisher et al., 2012). Predicting outbreak scenarios that involve a naïve host species and an emerging pathogen continues to be a global unsolved challenge (Bozzuto et al., 2020, and references therein).

A worrisome and truly global example that fits the constellation of naïve populations, introduced carriers and a lethal pathogen are native freshwater crayfish species confronted with invasive crayfish and the water mold *Aphanomyces astaci*. Crayfish are decapod crustaceans belonging to the families *Astacidae, Cambaridae* and *Parastacidae* (Kawai & Crandall, 2016). They are of central importance for freshwater ecosystems for their control of trophic webs and their role as ecosystem engineers (Reynolds et al., 2013). The native global distribution of freshwater crayfish includes North- and South-America, Eurasia, Australia and New Zealand, and Madagascar (Kawai & Crandall, 2016). Invasive crayfish species – mainly by human-mediated transportation both as food organism and ornamental species – are reported on all continents except Antarctica and cause enormous ecological and economic impacts (Kouba et al., 2022; Vaeßen & Hollert, 2015). These impacts are caused by a combination of direct ecological effects (competition with native species) and indirect effects (pathogen transmission to resident naïve native species; Gherardi, 2007).

Native crayfish species outside of North America are under pressure from the water mold *A. astaci* which is the etiological agent of a fatal disease commonly referred to as crayfish plague (Holdich et al., 2009). The pathogen likely originated in North America, whose crayfish species are rarely fatally affected and co-evolved with the pathogen (Martín-Torrijos et al., 2021). *A. astaci* is not the only pathogen affecting non-North American crayfish populations, but it is the pathogen with the most devastating impact (Jernelöv, 2017; Longshaw, 2011). Indeed, it has been slowly and steadily spreading globally, mainly due to human activities (Kawai & Crandall, 2016; Kouba et al., 2022; Longshaw, 2011; Reynolds et al., 2013; Vaeßen & Hollert, 2015). Native European crayfish species, which have not co-evolved with *A. astaci*, mount an inefficient immune response, and infection leads almost always to population-scale die-off events and local extinctions (Svoboda et al., 2017). Despite the fatal impact of *A. astaci* on native populations, crayfish populations are usually isolated enough from one another to make a natural invasion (i.e., not mediated by humans) of the pathogen less likely. Nevertheless, due to the continued presence of invasive (carrier) species, the pathogen has been persisting in Europe for ∼150 years and continues being an existential threat to the remaining native crayfish populations.

The pathogen is mainly transmitted through water by biflagellate spores called zoospores which are chemotactically attracted to crayfish tissue (Koivu-Jolma et al., 2023; Olson et al., 1984). Once zoospores reach a susceptible host, they form cysts which germinate and invade the exoskeleton (Nyhlén and Unestam, 1980). The pathogen is highly dependent on crayfish and cannot survive more than two weeks in distilled water without crayfish tissue (Unestam, 1966).

Management of crayfish plague in natural water bodies is difficult (Theissinger et al., 2021). Once an outbreak occurs, the pathogen cannot be eliminated without removing possible susceptible hosts or without harming other animal species, e.g. through chemicals, thus creating biodiversity damage. Disease management is therefore mainly based on the removal of invading non-native individuals, and on preventing involuntary animal and anthropogenically mediated pathogen transport from affected to non-affected water bodies (N’Guyen et al., 2018). This prevention requires reliable detection of the pathogen, of non-native carrier species, and targeted reduction of contacts between native and non-native carrier species. Given the large resource requirements for both preventative and mitigation management, and considering the fragmented landscape of suitable habitats, evidence-based prioritization of management areas is essential for successful long-term protection of selected native crayfish populations.

A promising theoretical background for wildlife disease-related mitigation plans is epidemiological theory. Central metrics like the basic reproduction number *R*_0_ have been developed a century ago and have since been used to understand and manage human and wildlife disease epidemics (Joseph et al., 2013; Keeling & Rohani, 2008). Surprisingly, research on crayfish diseases to this day has been characterized by a glaring absence of epidemiological concepts to guide mitigation actions. To our knowledge, a single recent study has investigated the epidemiological dynamics of host populations affected by crayfish plague (Koivu-Jolma et al., 2023). This is astonishing, given the long history of crayfish plague and crayfish introductions in Europe (Jernelöv, 2017), and the classification of crayfish plague as reportable disease in several European countries.

Epidemiological theory is primarily concerned with understanding conditions under which a host population will suffer a pathogen-caused epidemic, and further with shedding light on the middle- and long-term consequences such as endemic prevalence. One fundamental metric is the basic reproduction number, *R*_0_, giving the number of secondary infections caused by one infected individual in a completely susceptible population. This metric has been successfully adapted to diverse epidemiological situations, including vector-borne pathogens or diseases with carrier states, just to name a fraction of the many possibilities. A leap forward came in the 1990s with the development of a method (next-generation matrix) to reflect host heterogeneities (Diekmann et al., 1990). The method can be applied to study within-host species differences affecting disease spread, but also to multi-host species communities where outbreak dynamics are strongly driven by among-host species differences in susceptibility. Another important aspect stressed by Bowers and Turner (1997), and by Roberts and Heesterbeek (2013, 2018, 2020) is not just considering epidemiological dynamics in isolation, but to include the ecological context using a so-called eco-epidemiological approach (Figure 1A). Such frameworks account for the fact that susceptible populations and pathogens do not live in isolation, and embedding them in an ecological context can accordingly impact predictions about the effective spread of the disease and about effective mitigation options (Cortez & Duffy, 2021, Fenton et al., 2015, Rudge et al., 2013, Wilber et al., 2020).

**Figure 1.**
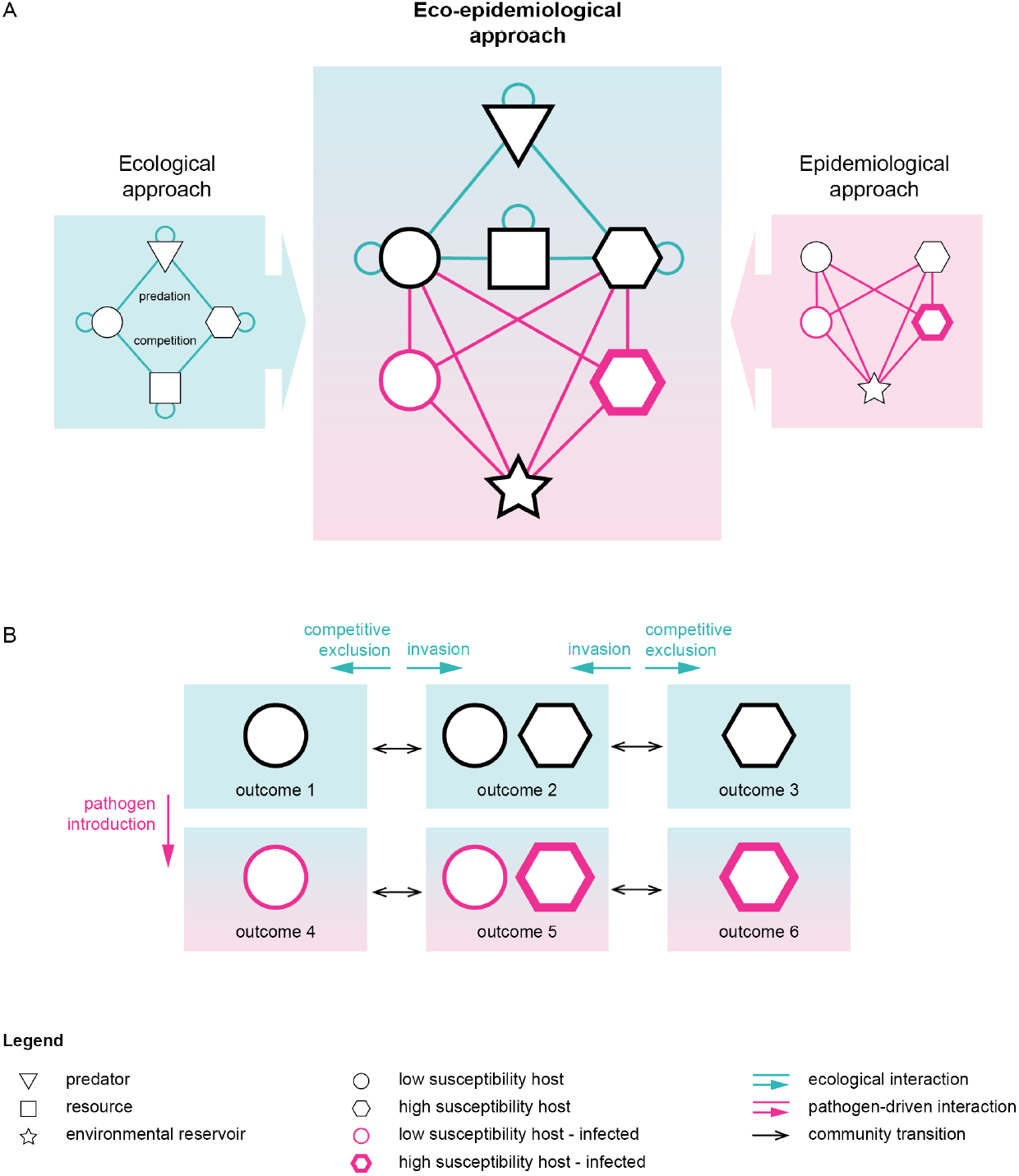
Principles of an eco-epidemiological modelling framework. **A –** Eco-epidemiological approaches combine knowledge about ecological interactions (cyan) with knowledge about disease-mediated effects (magenta). **B –** The community-level outcomes predicted by eco-epidemiological approaches therefore may differ from purely ecological or purely epidemiological approaches and have the potential to offer additional insights for preventative and mitigation actions.

The goal of our article is to present a flexible and generalizable eco-epidemiological framework to predict disease-related community outcomes involving an invasive pathogen (*A. astaci*) and host species with different susceptibilities (native and invasive freshwater crayfish) embedded in their ecological context. Our aim is to provide a tool to prioritize preventative and mitigation actions. We start by showing the consequences of considering multiple host and non-host species (with respect to a pathogen) for disease dynamics, presenting simple expressions of *R*_0_. Against this background, we then address the application of our findings to inform management, ranging from local single-host species epidemics to regional risk rankings spanning a variety of single- and multi-host species communities. We adopt a verbal, conceptual presentation in the main text, referring the reader to the Supporting Information for technical details where appropriate.

## Materials and Methods

The aim of our approach is to derive management-relevant insights from the ecological and epidemiological forces shaping the dynamics of crayfish communities confronted with crayfish plague. Even if the focus is on a community composed of two crayfish species that share the same habitat, additional biotic (e.g. predators) or environmental aspects (e.g. environmental pathogen reservoir) will drive the eco-epidemiological dynamic of the community.

### What constitutes a community?

A key characteristic of communities is the quality and quantity of the direct and indirect interactions among the factors that affect eco-epidemiological dynamics (Fig. 1-2). For example, is an interaction unilateral (e.g. crayfish as prey of a generalist predator) or bilateral (e.g. competition among two crayfish species)? Are there indirect interactions among crayfish species mediated by an a/biotic factor (e.g. infection through an environmental pathogen reservoir to which both species contribute)? In the absence of interactions, two crayfish species sharing the same habitat will display independent population dynamics. An eco-epidemiological model seeks to summarize relevant feedback mechanisms along both the ecological and the epidemiological axis.

### Ecological dynamics

Crayfish species are omnivorous, and therefore it makes little sense to include all possible prey species in a mathematical model. Instead, for a single species a simple model with density-regulated population growth will suffice in many cases (Supporting Information, section S2: eq. S1a). If two crayfish species share the same habitat and they compete for resources (i.e. direct interactions: compare Fig. 2A and 2B), both species-specific models should include competition terms: the higher a competitor’s density is, the lower the population growth rate of a focal species is expected to be (Supporting Information, section S2: eq. S1a-b). Although we focus on a two-species community – probably the most common form of multi-species crayfish community – the same approach carries over to three-species communities (May & Leonard, 1975). Crayfish are preyed upon by different predators like birds and fish, and by humans via fishing. These can be regarded as generalist predators, which renders the interaction unilateral: predation will lower a crayfish population’s density and growth, but the (long-term) dynamics of the predator population are largely unaffected by the density of a single prey species. In contrast to competition effects, the effect of generalist predators will enter the model of a focal species as their (density-independent) population growth rate reduced by a constant; the dynamics of a predator’s density therefore is not explicitly tracked through time. In the Supporting Information, section S2, we explain why fixing a predator’s density to a constant also works for cases where a crayfish species preys upon eggs or juvenile stages of its own predator. In sum, this set of mechanisms allows setting up a model mirroring the main factors affecting the dynamics of a crayfish community (Fig. 3; Supporting Information, section S2: eq. S3a-b). With this model in hand, the question of the possible outcomes over time can be investigated: what will affect coexistence or competitive exclusion (Fig. 1B)?

**Figure 2.**
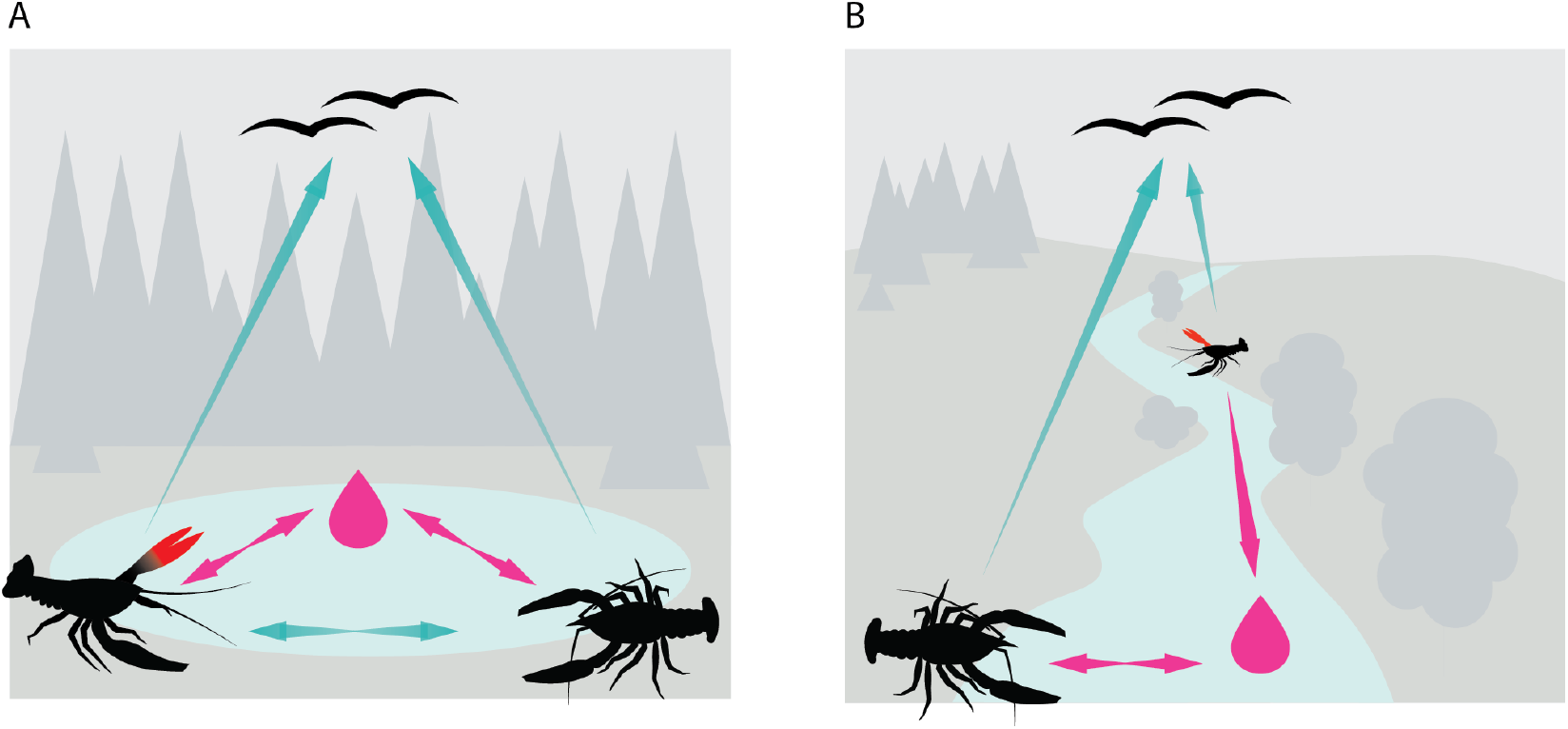
Illustration of communities in their habitat. **A –** Two crayfish species compete for resources in the same water body, are both preyed upon, and are affected by the indirect transmission of the pathogen *A. astaci* through spores via the water body (environmental reservoir) **B –** Two crayfish species inhabit the same water body (river) but at a ‘considerable distance’, so that they do not compete for resources, and the environmental pathogen reservoir of the down-river species is increased by the pathogen inflow from the up-river species, but not vice versa. Ecological interactions: cyan, epidemiological interactions: magenta.

**Figure 3.**
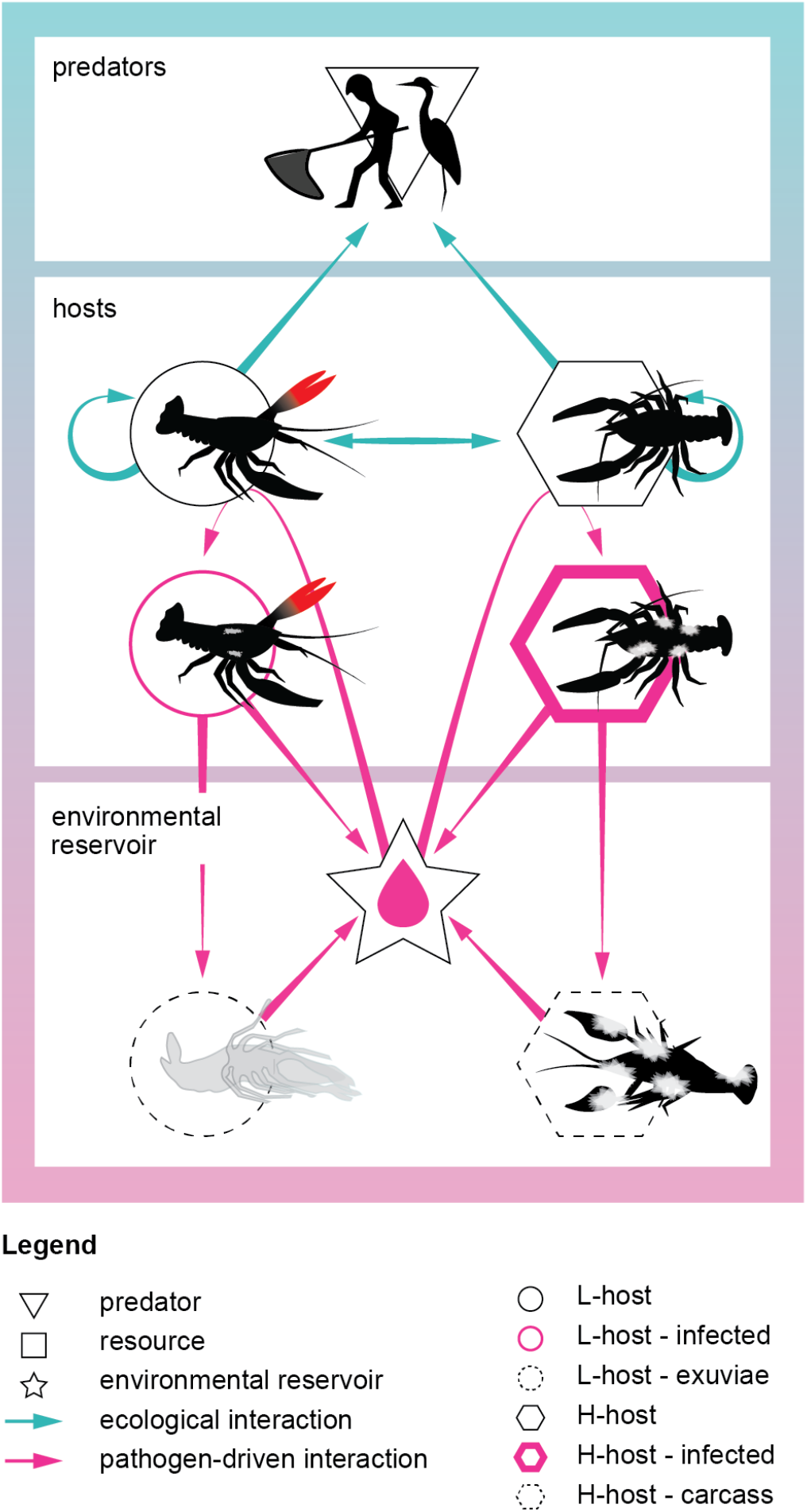
Illustration of eco-epidemiological model for a crayfish community. The figure shows the main ecological and epidemiological processes forming our eco-epidemiological model; for ease of visualization, loss-of-infectiousness rates (Table 1) are not shown. The underlying symbols (legend) are the same as in Fig. 1. Ecological interactions: cyan, epidemiological interactions: magenta.

### Epidemiological dynamics

Epidemiological models of a single host species and a single pathogen summarize, akin to the ecological approach described above, the relevant mechanisms driving disease-related population dynamics. A common approach is to divide the host population into disease-related compartments, for example susceptible or infected individuals (e.g. Keeling & Rohani, 2008). For every compartment, an equation describes how disease-related ‘entities’ (e.g. living or dead individuals, carcasses etc.) enter (e.g. through infection) or leave the compartment (e.g. through recovery or degradation; Supporting Information, section S3: eq. S5a-e). For this type of modelling approach, pathogen dynamics are usually not explicitly modelled: a pathogen is simply mirrored in infectivity- or virulence-related parameters, among others. For a community of multiple host species, each host population has its own set of epidemiological equations, but among-host species processes (e.g. transmission) are also accounted for. Finally, the single- or multi-host setting can be supplemented with additional equations, for example to describe an environmental pathogen reservoir. Fig. 3 depicts the main underlying mechanisms forming our eco-epidemiological model, which we verbally summarize below. Additional mechanisms not included in the model, such as direct among-host species transmission – and the reasoning to not consider them – are presented in the Supporting Information, section S3. Henceforth, the nomenclature is as follows: (i) host species naïve with respect to *A. astaci* (i.e., native European crayfish species) are called “high susceptibility host” (short: H-host); (ii) host species immune to *A. astaci* (i.e., North American crayfish species) are called “low susceptibility host” (short: L-host); (iii) carcasses and exuviae (moulted exoskeleton) that release pathogen spores, as well as pathogen spores present in the water, are called “environmental reservoir”. Finally, for illustrative purposes, we focus on a crayfish community composed of an H-host and an L-host (but see the *Results* and *Discussion* for more complex scenarios). All model variables and parameters are summarised in Table 1.

**Table 1.**
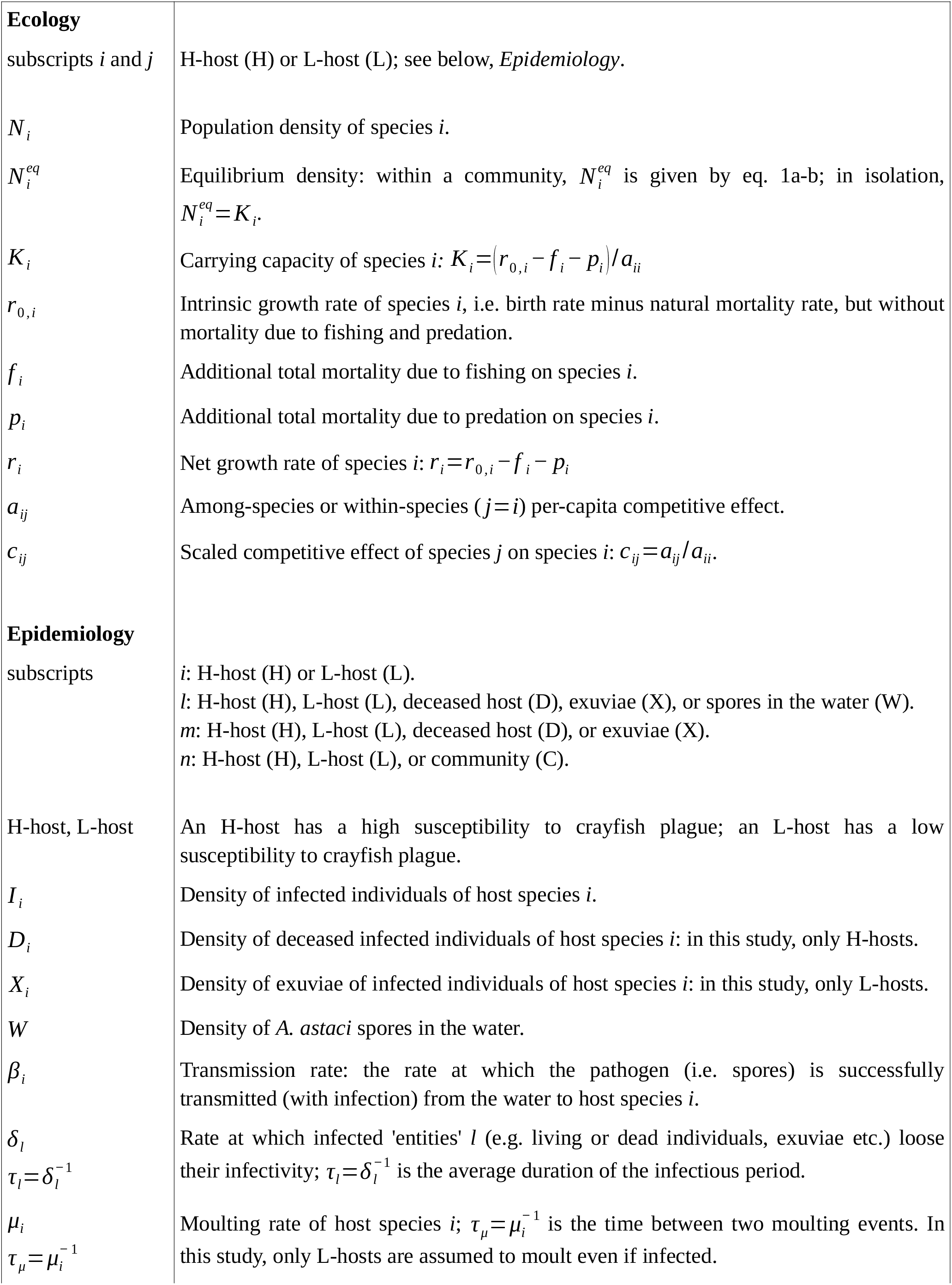
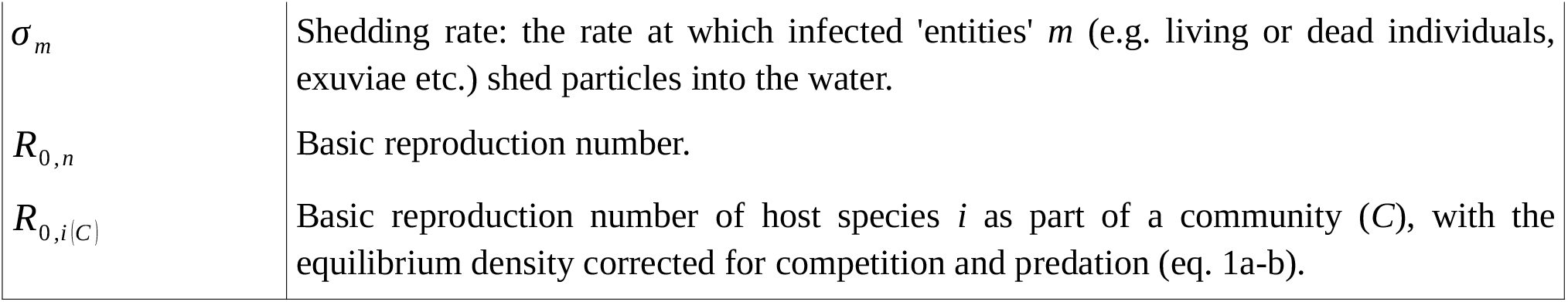
Summary of variables and parameters used in this study.

The main processes affecting the epidemiological dynamics are:

▪ Susceptible individuals of an H-host are infected by spores in the water, at a rate specific to the host species. Infected H-host individuals remain infectious for a certain period, after which they die.
▪ Susceptible individuals of an L-host are infected by spores in the water at a rate specific to the host species. Infected L-host individuals remain infectious for a certain period, after which they loose infectiousness.
▪ The bodies of deceased infected individuals of an H-host will continue shedding pathogen spores into the water for a certain period, after which they can be considered decomposed and not infectious anymore.
▪ The exuviae of infected individuals of an L-host (after moulting) will continue shedding pathogen spores into the water for a certain period, after which they can be considered decomposed and not infectious anymore.
▪ The density of pathogen spores in the water increases due to particle shedding of infected individuals of both host species, their carcasses (H-hosts) and exuviae (L-hosts). It decreases due to particle degradation and transportation.

In sum, compared to L-hosts the difference of H-hosts lies mainly in the disease-induced mortality (instead of loosing infectiousness alive) and their carcasses (instead of exuviae) continuing shedding pathogen spores into the water.

As with ecological models, epidemiological ones allow studying different aspects of the dynamics, for example which circumstances promote an outbreak (i.e. epidemic), or asking about the density of the different compartments once an endemic state is reached. Here, we focus the basic reproduction number, *R*_0_, because it gives important insights into initial outbreak dynamics, among others. For example, *R*_0_ = 1 acts a threshold with values above 1 indicating an outbreak to be expected. Moreover, the *R*_0_ value often correlates with the speed with which a pathogen spreads through a host population (Diekmann et al., 1990). We derive *R*_0_ expressions for H-hosts and L-hosts individually, as well as for the community as a whole. We then use these expressions to propose ways of guiding management and mitigation decisions. Given the already fragmentary ecological and epidemiological knowledge on crayfish and crayfish plague, we have refrained from additionally considering spatial dynamics in our framework, in particular since it has been shown for other host-pathogen systems that non-spatial *R*_0_ expressions are also valid to study outbreak dynamics in different spatio-temporal situations (Zhang et al., 2016).

### Eco-epidemiological dynamics

The coupling of ecological and epidemiological dynamical processes unveils aspects and outcomes that either of them would not detect individually. Simply put, an eco-epidemiological approach allows studying the conditions for stable coexistence (or lack thereof) between two or more host species, in the absence or presence of a pathogen, accounting for all relevant abiotic and biotic ecological and epidemiological factors affecting these dynamics (Fig. 1B). Because we focus our attention on the conditions enabling an outbreak to occur, with *R*_0_ as our metric, we first study a community in a pathogen-free environment, followed by investigating the conditions for this community – when confronted with crayfish plague – enabling an epidemic. We do not focus on a potential endemic state (Supporting Information, section S1) because previous research suggests that once a naïve crayfish population is affected by the disease, the likely outcome is local extinction (Alderman et al., 1987; Martín-Torrijos et al., 2017; Schrimpf et al., 2013).

### Deriving management-relevant insights from a non-parametrized model

Management-relevant insights can be derived by parameterising eco-epidemiological models and related metrics, like ecological equilibrium densities or *R*_0_, and to connect potential preventative and mitigation actions to numerical changes in these metrics. This step is usually very challenging in wildlife disease management (McCallum, 2016), and at the current state of knowledge a full parameterisation is not feasible for most crayfish species and communities. However, there are alternative ways to gain insights that circumvent the need for absolute parameter values.

First, the metrics of interest can be inspected qualitatively. For example, as we will show the *R*_0_ value of a crayfish community is the sum of the community’s single-host species *R*_0_ values, adjusted for competition and predation, and this could already coarsely help comparing and ranking different communities for a management prioritisation.

Second, a related semi-quantitative approach is to perform a sensitivity analysis (Supporting Information, section S4). In a nutshell, for parameters or variables of interest (potentially influenceable by management actions), a sensitivity analysis indicates how a slightly changed parameter or variable value will affect a metric of interest like *R*_0_ (e.g. Canessa et al. 2019; Caswell, 2019). Although such sensitivity analysis will be most efficient for fully parameterised models, simple mathematical techniques already allow coarsely ranking management actions in terms of their (expected) effectiveness. In practice, however, budgetary effectiveness will also play a crucial role, and therefore a final ranking of available mitigation actions will also be based on (long-term) resources needed per action (not pursued further in this study). Note that, based on the mathematical approach underlying a sensitivity analysis, if the management-induced proportional changes are small (sic!), then the results could also be used additively to study combinations of mitigation actions (Caswell, 2019).

Third, to compare several communities to one another in terms of disease outbreak risk we propose to build *R*_0_ ratios (see also Bielby et al. 2021; Fenton et al. 2015), and this approach will be especially useful for a regional ranking within a prioritisation process. The ratio of two *R*_0_ values can be mathematically manipulated so that all analogous host species-specific parameters also appear as ratios in the *R*_0_ ratio. The use of an *R*_0_ ratio simplifies the comparison of two communities: it is often feasible to gauge a parameter value of one species relative to a second one. For example, it may turn out that a parameter related to transmission is expected to be 2 or more times higher in one species than in the other species. These relative estimates can often be given with confidence by environmental managers, stakeholders or researchers, and abolish the need for precise parameter estimates when using *R*_0_ ratios for management prioritization.

## Results

### Ecological dynamics

In a pathogen-free environment, the ecological dynamics in the presence of generalist predators (Fig. 3) allow competitive exclusion or coexistence as possible long-term outcomes (Supporting Information, section S2) for a crayfish community. We focus on coexistence (Schrimpf et al., 2013), for which the equilibrium densities of an H-host 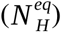 and an L-host 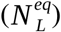 forming a community are

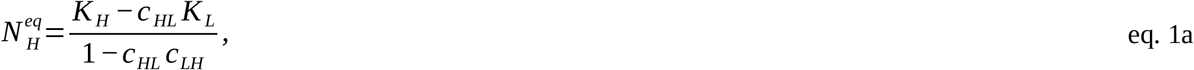

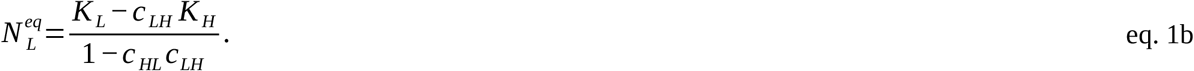

In these two expressions, *K* _*H*_ and *K* _*L*_ are the carrying capacities of the two host species in isolation, and all *c*_*ij*_ are competition-related parameters (Table 1); eq. 1a-b are well known results in mathematical ecology (e.g. Kot, 2001). Parameterisation of these expressions will be challenging in practice. Nonetheless, these expressions offer general insights useful to guide preventative and mitigation actions. First and expectedly, strong competition from an L-host (a high *c* _*HL*_ value) will lower the equilibrium density of an H-host (albeit in a nonlinear way), but such decrease is opposed by the competition of the H-host towards the L-host (*c* _*LH*_). Second, with respect to the carrying capacities it may be feasible to estimate them, or to use literature values. But a general insight can nonetheless be gained from the definition of a single-host species carrying capacity (Table 1; Supporting Information, section S2), *K*_*i*_ =(*r*_0,*i*_ − *f* _*i*_ − *p*_*i*_)/ *a*_*ii*_. Thus, the equilibrium densities of two coexisting species (eq. 1a-b) depend linearly on host-specific mortality due to fishing (*f* _*i*_) and predation (*p*_*i*_), and this insight may help better understand disease outbreak dynamics (see below, sub-section *Preparedness and outbreak emergency*).

### Epidemiological dynamics

Given our focus on crayfish communities inhabiting a pathogen-free environment and confronted with the pathogen *A. astaci*, a main interest in constructing an eco-epidemiological framework lies in the derivation of basic reproduction numbers for single host species and communities. Our epidemiology-related assumptions (Fig. 3) lead to a simple and handy result (Fig. 4; Supporting Information, section S3, eq. S10): the community-level basic reproduction number, *R*_0,*C*_, is the sum of the host species-specific basic reproduction numbers, with host densities adjusted for competition and predation effects. For example, for a community composed of an H-host and an L-host, *R*_0,*C*_ = *R*_0, *H* (*C*)_ +*R*_0,*L* (*C*)_; here, subscripts *H*(*C*) and *L*(*C*) (see also Table 1) mean that the respective host equilibrium densities include competitive and predatory effects (eq. 1a-b). This simple relationship results from the realistic assumption that the main pathogen transmission route happens indirectly among host individuals via the water (instead of direct contact transmission), when the two host populations have an overlapping habitat and therefore both contribute to and take up spores from a shared environmental reservoir (Fig. 2A).

**Figure 4.**
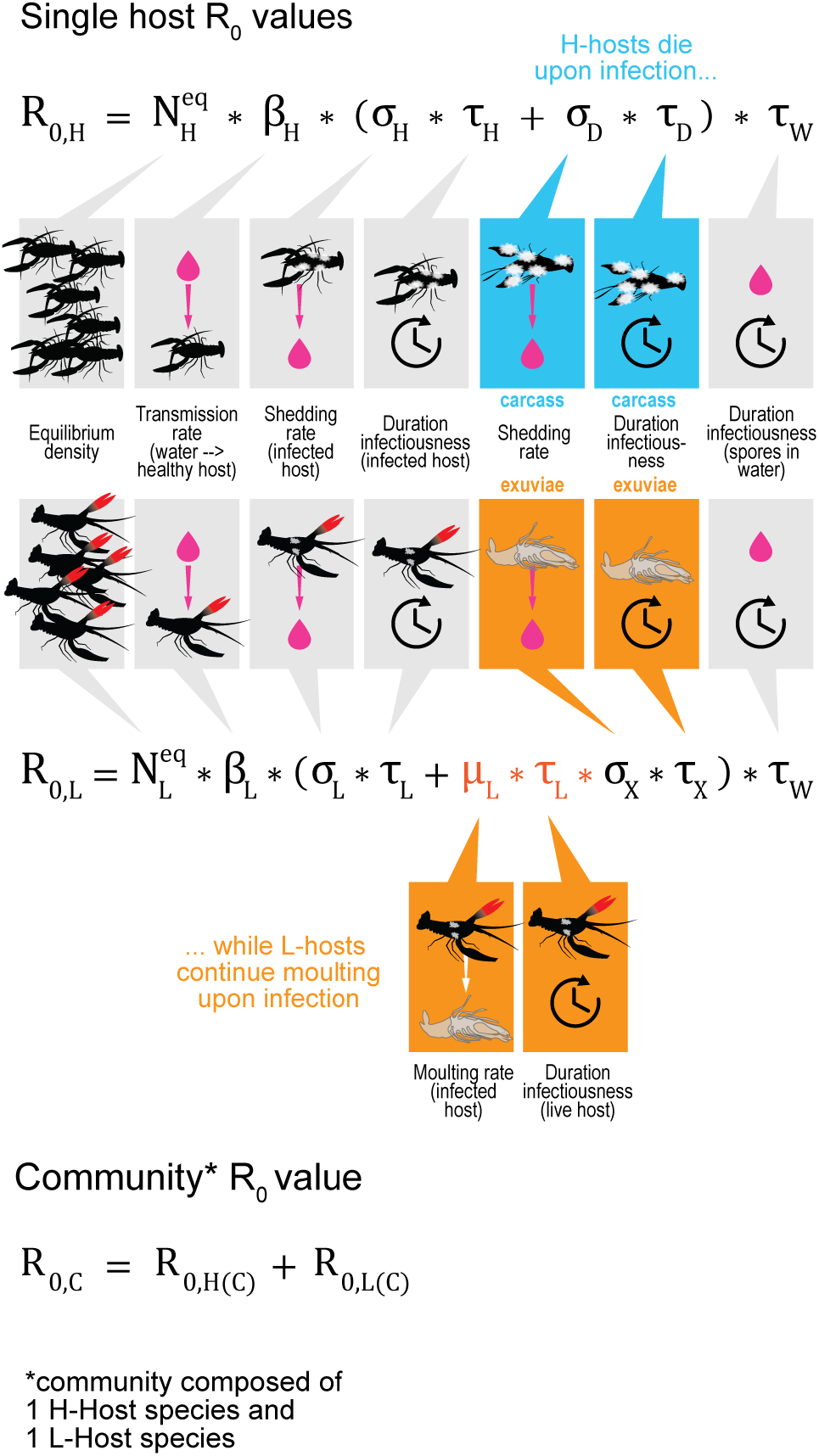
Basic reproduction numbers for single host species and communities. The top figure part shows the single-host species *R*_0_ expressions (H-host, L-host), incl. a graphical explanation of the constituting parameters (see also Table 1). If a host species has no competitors in its habitat, 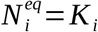, otherwise 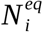 is given by eq. 1a-b. The bottom figure part shows that a crayfish community’s *R*_0_ value (*R*_0,C_) is the sum of the community’s single-host species adjusted for competition and predation (*R*_0,H(C)_ and *R*_0,L(C)_).

### Eco-epidemiological dynamics

The single-host species and community *R*_0_ expressions shown in Fig. 4 evidence the dependence of the expected likelihood and severity of an outbreak on both the ecological and epidemiological context. On the ecological side, an *R*_0_ value will increase with increased equilibrium density which depends on life-history characteristics, competition strength and trophic interactions (eq. 1a-b, Fig. 3). On the epidemiological side, changes in *R*_0_ depend on characteristics of the host-pathogen interaction (e.g. virulence), but also on indirect transmission pathways via the environment (Fig. 3-4). Thus, an eco-epidemiological framework not only allows deriving the outbreak-related metric *R*_0_, it moreover allows deriving management-relevant insights by inspecting *R*_0_ expressions.

### Preparedness and outbreak emergency

A common way for *R*_0_ to inform potential mitigation actions – in terms of preparedness, preventing an outbreak, or curbing an epidemic curve – is to understand what actions (which influence variables and parameters) can keep or drive *R*_0_ below or near the outbreak threshold *R*_0_ = 1. Despite its high utility, this approach unfortunately requires numerical evaluation of *R*_0_ in-/excluding actions. Two less stringent, but nonetheless informative approaches are to understand (i) which potential actions could realistically drive desired changes in parameters and variables and ultimately in *R*_0_, and (ii) which parameter or variable change(s) will entail the most marked change in *R*_0_.

As for the first question, for example for an H-host inhabiting a habitat alone (i.e. without additional crayfish species present) the expression for *R*_0_ suggests four actions to lower its value: (i) (temporarily) reduce host population density 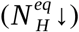 (ii) selectively remove infected individuals (*τ*_*H*_ ↓); (iii) remove dead animals (*τ*_*D*_ ↓); (iv) reduce the spores’ persistence in the water (e.g. by chemical and/or physical water treatment: *τ*_*W*_ ↓). Action (i), however, can represent a dilemma for managers. For a vulnerable/endangered H-host, decreasing its density is not desired for conservation-related reasons (at least in the long-term), but a sensibly decreased density could potentially help prevent an outbreak (*R*_0_ < 1). Things change if such H-host forms a community with an L-host: in this case community *R*_0_ could be decreased by reducing the equilibrium density of the L-host (eq. 1a), for example via targeted increased and sustained fishing effort (see below).

To address the second question – which parameter or variable change(s) will entail the most marked change in *R*_0_ – a sensitivity analysis can be performed. The goal is to ask how a slightly changed parameter or variable value will affect a metric of interest, in the present case *R*_0_, with the aim of ranking such management actions in terms of their efficiency. For a single H-host, elasticity expressions show that an equal proportional decrease in 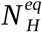or *τ*_*W*_ will entail the same proportional change in *R*_0_; in comparison, proportional changes in *τ*_*H*_ and *τ*_*D*_ are less effective to achieve the same change in *R*_0_ (Supporting Information, section S4: eq. S13a-c).

To use a sensitivity analysis for a crayfish community, two situations might be considered separately. One is an “emergency” situation, where mitigation actions would be deployed not long before an outbreak is expected to happen, for example because of recently observed outbreaks in adjacent water bodies (e.g. Canessa et al., 2019). Here, the time scale is expected not to be long enough for ecological processes like competition to react to the manipulation of host species densities as one potential action (see above). Thus, *R*_0,*C*_ (Fig. 4) can be used by assuming that both host species densities are independent of each other (as opposed to eq. 1a-b). The parameters that potentially can be influenced by mitigation actions are thus 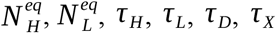. The respective elasticity expressions (Supporting Information, section S4: eq. S14a-g) show that reducing the persistence of spores in the water appears to be the most efficient option. Reducing host species densities in a community is now less efficient than the former option, compared to a single host population in isolation (see above). In addition, the elasticity expressions allow ranking other options more finely: for example, for the L-host we have 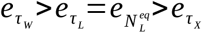. A full ranking, nonetheless, will require a full parametrization.

As for the second situation with respect to a crayfish community to be managed, we assume that some management actions may be applied over a longer time period, without an expected incumbent outbreak. In fact, for such situation we have different relevant time scales: host species densities can be decreased for example by fishing – in a sustained way – over a longer period of time if appropriate (see below) and in the absence of an outbreak, while other actions are necessarily tied to an outbreak (e.g. reducing durations of the infectious period). Because the manipulation of host densities could happen over a longer period of time, in contrast to an emergency situation (see above) competitive effects are now expected to react to density manipulations. Thus, in addition to the duration of infectious periods already analyzed for an emergency situation (*τ*_*H*_, *τ*_*L*_, *τ*_*D*_, *τ*_*X*_, *τ*_*W*_), we now add the carrying capacity of the L-host to this list, *K* _*L*_ : the intent here is to reduce the density of the invasive host species in a sustained way, and therefore to indirectly increase the density of the native host species (eq. 1a-b). The elasticity values and relative ranking for the duration of the infectious periods are the same as for the emergency situation described above. The inspection of the elasticity expression for *K* _*L*_ (Supporting Information, section S4: eq. S16) reveals an important caveat with respect to a long-term decrease of the density of an L-host as a management action: despite the good intention to indirectly increase the long-term density of a susceptible and potentially endangered H-host (eq. 1a), density manipulation could backfire in terms of outbreak risk, i.e. a desired *reduction* in *K* _*L*_ could unintentionally lead to an *increase* of *R*_0, *C*_. Whether or not this might be the case for a specific crayfish community can only be answered after a parametrization. Nonetheless, the elasticity expression for *K* _*L*_ suggests that the backfiring is a real possibility. Moreover, crayfish communities with host species attaining a similar carrying capacity in isolation (*K*_*i*_) will increase the likelihood of a higher *R*_0,*C*_ value when preemptively reducing the long-term density of an invasive L-host (Supporting Information, section S4).

### Regional risk ranking

Given limited resources for conservation actions (Walls, 2018), a sensible way of planning resource allocation to single conservation projects is to first gain a regional overview in terms of a prioritization. Hereto, we propose the comparison of single-host species and community-level *R*_0_ expressions for a (larger) region, potentially containing different single-host species and communities composed of different combinations of host species (Fig. 5A). The idea is simple: start by choosing a single ‘baseline’ host species – in central Europe for example the native highly susceptible noble crayfish (*Astacus astacus*) – and then conduct pairwise comparisons with other single host species or communities by building *R*_0_ ratios (Fig. 4). These ratios will reflect the relative likelihood of an outbreak for a given host species or community, compared to the ‘baseline’ host species (Box 1). Therefore, these ratios allow coarsely ranking regional crayfish-inhabited water bodies in terms of conservation priorities: values of ratios exceeding 1.0 will indicate a higher priority with respect to the ‘baseline’ species (Fig. 5B).

**Figure 5.**
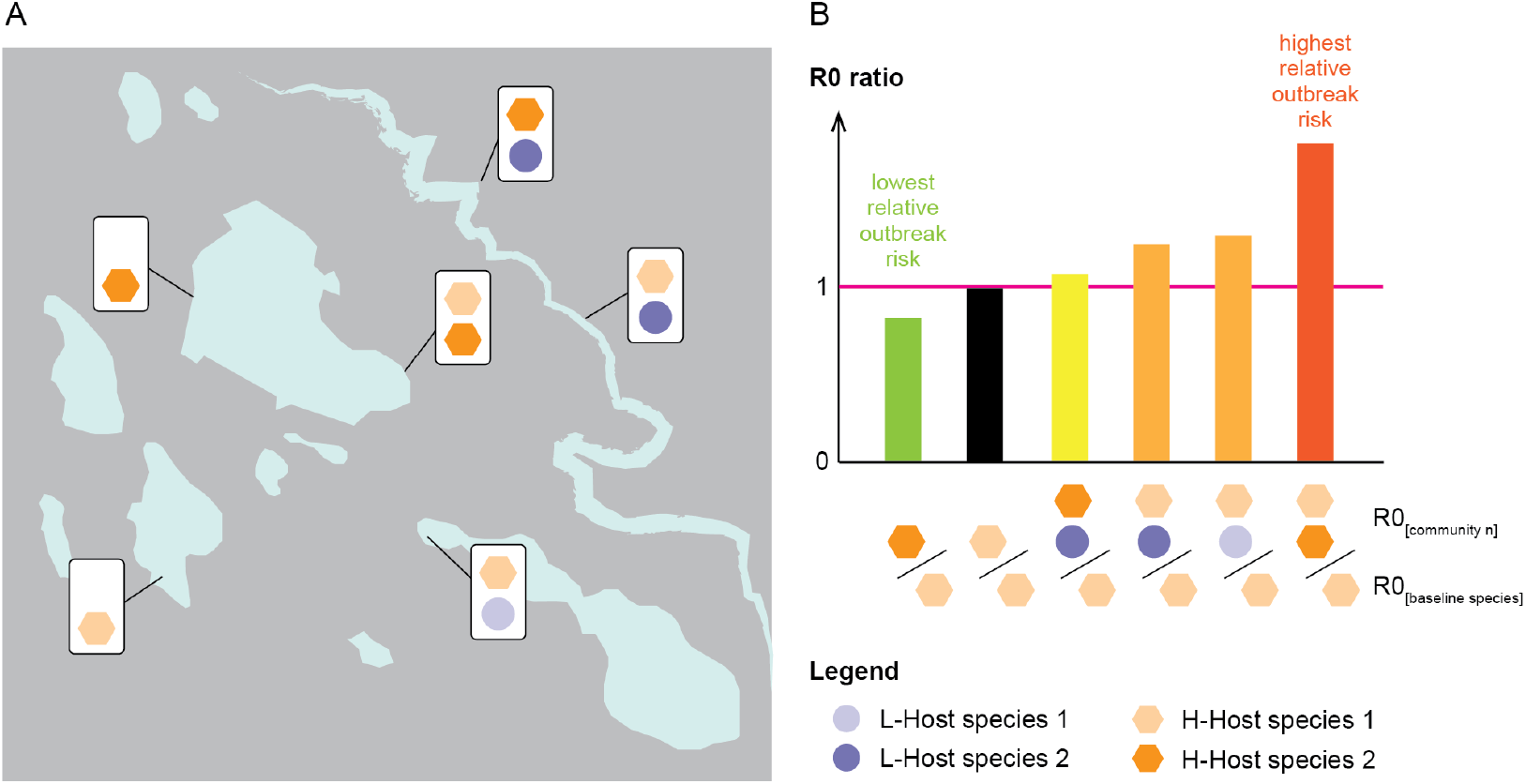
Illustrative regional risk ranking of six crayfish communities. **A –** An illustrative region with different water bodies inhabited by different crayfish communities (single and multiple species). **B –** The bar diagram gives the *R*_0_ ratio values – in ascending order, i.e. relative outbreak risk – for all communities shown in panel A with respect to the ‘baseline’ H-Host species 1.

#### Box 1.

**Illustrative risk ranking**

To illustrate how a regional risk ranking based on single-host species and community-level *R*_0_ expressions can be achieved, here we theoretically compare *R*_0_ of a community composed of an H-host and an L-host (*R*_0,*C*_ = *R*_0, *H*(*C*)_ +*R*_0, *L*(*C*)_; Fig. 4) to *R*_0_ of the same H-host in isolation (*R*_0,*H*_). Hereto, we inspect the ratio

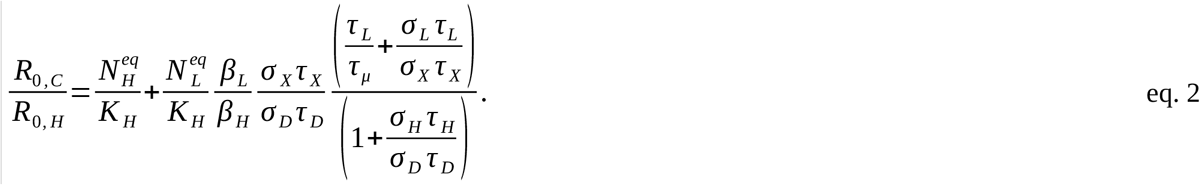

Eq. 2 is a rearranged version of the *R*_0_ ratio based on host-specific *R*_0_ expressions presented in Fig. 4 (Supporting Information, section S4: eq. S17-18), with the aim of presenting all parameters and variables as ratios. Thus, to coarsely gauge the value of an *R*_0_ ratio, the task becomes to understand whether and by how much a parameter or variable of one host species differs from the same parameter or variable of the second host species in a relative way. For example, the transmission of spores (and consequently of the disease) from the water to the hosts might be expected to be comparable, thus *β* _*L*_ / *β* _*H*_ ≈ 1.0. For eq. 2 we assume that the abiotic habitat is comparable, and therefore the spores in the water degrade at the same rate, which then cancels out when building the ratio. Eq. 2 could be further simplified if, for example, it is assumed that the shedding rate of an infected L-host is very low (*σ* _*L*_ ≈ 0.0); L-hosts in such case would only contribute spores to the environment through their infected exuviae. For the species constellation used to derive eq. 2, a further simplification (i.e. an approximation) could be to set 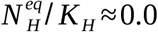 the reasoning is that the equilibrium density of an H-host in the presence of an invasive competitor (L-host) can be expected to be (much) lower than in isolation (e.g. Westman & Savolainen, 2001), thus this ratio may be expected to be (much) smaller than 1.0 and contribute little to the overall ratio. The approach of building a ratio can be used for comparing any two host species and/or communities. For comparisons of two host species in isolation (i.e. without competitors in their respective water body), the ratio is similar to the second summand in eq. 2, where now (i) both equilibrium densities are carrying capacities (Table 1), and (ii) the overall ratio should potentially also include a ratio for the persistence time of spores in the water (*τ*_*W*_) in case the compared host species live in contrasting environments with differential effects on *τ*_*W*_. As a second example, for comparisons where a community does not include the ‘baseline’ species, both summands in the overall ratio are similar to the second summand in eq. 2. For instance, if the community to be compared to the ‘baseline’ host species (H-host #1, say) is composed of H-host #2 and an L-host (Fig. 5B), then the ratio becomes *R*_0,*C*_ / *R*_0,*H*1_ =*R*_0,*H*2(*C*)_ / *R*_0*H*1_ +*R*_0,*L*(*C*)_ / *R*_0,*H*1_.

## Discussion

We have presented a general and flexible eco-epidemiological framework to study and aid management of the biodiversity “triad of death” (native species, invasive species, and invasive pathogen) applied to crayfish communities confronted with crayfish plague. Motivated by the glaring gap of quantitative approaches for such communities in the literature, our aim to inform conservation management was twofold. First, the construction of any quantitative eco-epidemiological model forces researchers and practitioners to make relevant known factors and mechanisms affecting ecological and epidemiological processes explicit, allowing a sound summary of crucial factors. Second, we demonstrate how derived metrics, especially the basic reproduction number (*R*_0_), can be used as a starting point to guide mitigation and prevention at the local and regional scale, for single populations and communities of crayfish host species, even in the absence of a comprehensive parameterisation.

We have started with ecological aspects of a crayfish community. Besides regional differences to be expected in different habitats and biogeographic regions, we show that crayfish communities are mainly characterized by life-history characteristics, competitive interactions and interactions with generalist predators preying upon crayfish, including exploitation by fishing. Population densities of crayfish hosts forming a community will directly affect disease outbreak dynamics (Fig. 4). These dynamics will be additionally affected by pathogen transmission routes and by host-pathogen interactions. Environmental transmission via the water can be considered the most relevant transmission route for crayfish (Fig. 3) (Koivu-Jolma et al., 2023), and we demonstrate that this simplifies *R*_0_ expressions at the community level: community-level *R*_0_ is the sum of single-host species *R*_0_ values, with host densities adjusted for competition and predation (see also Fenton et al., 2015, for similar results). One corollary of such simplification is that the framework can be used and analysed for different kinds of communities with ease in terms of the number of host species and their susceptibility to crayfish plague. Furthermore, we expect the framework – or adaptations of it – to be useful for any aquatic host-pathogen system where the main pathogen transmission route is through water instead of direct contact among host individuals (e.g. Jenkins et al., 2021).

As for many wildlife diseases (McCallum, 2016), relevant knowledge for model parametrization is scarce for crayfish plague, which is a challenge regarding the utility of quantitative eco-epidemiological approaches beyond general insights. Nonetheless, the framework helps evidencing knowledge gaps when it comes to parametrization: what ecological or epidemiological aspects have so far not been researched at all? What crucial eco-epidemiological processes are amenable to controlled studies and should be pushed and prioritised? To prevent for example a local single-host species outbreak, *R*_0_ is mainly useful to inform action planning if parameter values are available so as to relate the effectiveness of actions to the outbreak threshold *R*_0_ = 1. Fortunately, decision-making as a discipline offers many practical tools to tackle parameter uncertainty (Canessa et al., 2018). In addition, a sensitivity analysis can already help coarsely ranking single mitigation actions in terms of their expected efficiency to drive changes in *R*_0_. Furthermore, if expressions of *R*_0_ are used as ratios – as we have proposed for a regional risk ranking – quantitative thinking is facilitated by expert knowledge (ecologists, epidemiologists, veterinarians etc.) because now the focus is on relative parameter values rather than absolute ones. Assessing the relative magnitude of differences among two populations or communities thus helps inform the derivation of (regional) intervention options (Fig. 5).

In summary, the combination of invasive species and invasive pathogens, as exemplified by crayfish plague, is a particularly difficult biodiversity threat to manage (Theissinger et al., 2021). Mitigation actions to curb an unfolding disease outbreak in the wild have limited success rates, especially in settings that include the introduction of novel diseases into habitats with naïve host species (Bozzuto et al., 2020, and references therein). In parallel, resource limitations are common in biodiversity conservation, which necessitates the use of data-driven, focused, proactive and cost-saving actions (Walls, 2018). In this context, an increasing number of projects concerned with wildlife diseases include epidemiological theory to more effectively inform management decisions and improve interventions (Joseph et al., 2013). We hope that our proposed framework, or adapted versions of it, will thus be useful for the development of focused, realistic, and effective conservation interventions at the interface of ecological and epidemiological biodiversity threats.

## Supporting information

SupportingInformation

## Acknowledgements

We thank Raphael Krieg and Armin Zenker from the Koordinationsstelle Flusskrebse Schweiz for insightful crayfish-related discussions. We are also grateful to the anonymous reviewers and colleagues for their constructive feedback and helpful suggestions on improving the manuscript. This study has been made possible thanks to the financial support of the Swiss Federal Office for the Environment (grant no. 06.0121.PZ/0304) and the Swiss Federal Food Safety and Veterinary Office (grant no. 1.21.02).

## Availability of data

The article does not present any data.

## Authors contributions

All authors conceived and discussed the ideas and designed methodology; Claudio Bozzuto provided all quantitative models and performed their analysis; Simone R.R. Pisano and Heike Schmidt-Posthaus provided crayfish-related ecological and epidemiological information; Irene Adrian-Kalchhauser designed all figures; Claudio Bozzuto led the writing of the manuscript; all authors contributed critically to the drafts and gave final approval for publication.

