## SupportingInformation for "Towards an eco-epidemiological framework for managing freshwater crayfish communities confronted with crayfish plague"

### Supporting Information

#### Towards an eco-epidemiological framework for managing freshwater crayfish communities confronted with crayfish plague

Claudio Bozzuto[1], Heike Schmidt-Posthaus[2], Irene Adrian-Kalchhauser[2], Simone Roberto Rolando Pisano[2 *]

[1] Wildlife Analysis GmbH, Oetlisbergstrasse 38, 8053 Zürich, Switzerland.

[2] Institute for Fish and Wildlife Health, Vetsuisse Faculty, University of Bern, Länggassstrasse 122, 3001 Bern, Switzerland.

This PDF file contains:

- Additional methods information:
  - S1. Eco-epidemiological dynamics: overview
  - S2. Ecological dynamics
  - S3. Epidemiological dynamics
  - S4. Basic reproduction numbers as a basis for management
  - References Supporting Information

### Additional methods information

#### S1. Eco-epidemiological dynamics: an overview

The basic idea of an eco-epidemiological modelling approach is to construct – and subsequently analyse – a mathematical model that encompasses both ecological and epidemiological dynamics. For communities composed of several (host) species, there is a rich and established literature on ecological community dynamics (e.g. Kot, 2001; Turchin, 2003) and on epidemiological community dynamics (e.g. Diekmann et al., 1990; Keeling & Rohani, 2008). The combination of ecological and epidemiological dynamics within a single community model is less frequently done (cf. Bowers & Turner, 1997); but see well-established single-host models including demographic elements (Keeling & Rohani, 2008, and references therein).

In a series of articles, Roberts and Heesterbeek (2013, 2018, 2020) recently showed in detail how the concomitant analysis of ecological and epidemiological community dynamics will broaden the spectrum of community outcomes to be expected. For an eco-epidemiological model, an ecological sub-model and an epidemiological sub-model are connected. The link is a single or several population/s that act as a host to one or more pathogen/s. Despite this link, each sub-model can contain other species that only indirectly influence the host-pathogen dynamics. For example, the ecological sub-model may contain equations describing the dynamics of competitive or predatory species which, however, are not hosts to the pathogen but interact with the focal host species. Similarly, the epidemiological sub-model may contain equations describing the effect of abiotic or biotic factors affecting disease spread, but not directly affecting the ecological dynamics of a host.

An eco-epidemiological community analysis will focus on at least three questions (Fig. 1B in the main text). First, what kind of dynamics will a community of pathogen-related hosts and other species like predators or resources display over time, in the absence of one or several pathogen(s)? Second, what are the conditions for one or several pathogen(s) to elicit an epidemic? Third, can the system composed of host species, non-host species and pathogen(s) lead to an endemic state? An illuminating and recommendable worked-through example has been published by Roberts and Heesterbeek (2021), where the authors study the eco-epidemiological context of communities of red and grey squirrels in the UK when confronted with the squirrel pox virus.

In the present article, we focus on the first and second question. One reason is that previous research suggests that once a native, high-susceptibility crayfish population is affected by the disease, the likely outcome is local extinction (Alderman et al., 1987; Martín-Torrijos et al., 2017; Schrimpf et al., 2013). Therefore, we focus on a disease-free community to understand a) how ecological factors affect the (longterm) equilibrium densities of the crayfish species, and b) what happens to this community at the beginning of the introduction of a pathogen, in our case crayfish plague. A second and more technical reason is that the analysis of an endemic state will require model details to be defined (e.g. nonlinear uptake rate of spores from the water), including additional parameters, which are unlikely to be know currently, but which to a first approximation are not necessary to understand outbreak dynamics. Furthermore, studying the eco-epidemiological endemic state of a community will usually call for numerical approaches, requiring parameter values which are seldom known for crayfish communities and crayfish plague.

#### S2. Ecological dynamics

To analyse the ecological dynamics of a community in the absence of a pathogen, we follow Bowers & Turner (1997) and Roberts & Heesterbeek (2013) and use a Lotka-Volterra framework. Specifically, we consider (i) density-independent within-population growth, (ii) within-population competition for resources, (ii) among-population competition for resources, and (iii) effects of predation (incl. fishing); see also Fig. 1. Disregarding predation for now, the population growth rates of an H-host and an L-host population forming a two-species community can be expressed using differential equations as

$$\dot{N}_H = r_{0,H} N_H - \left(a_{HH} N_H + a_{HL} N_L\right) N_H, \quad \text{eq. S1a}$$

$$\dot{N}_L = r_{0,L} N_L - \left(a_{LL} N_L + a_{LH} N_H\right) N_L, \quad \text{eq. S1b}$$

where a dot above a variable denotes a time derivative; for notational simplicity we omit the time dependence of all ecological and epidemiological variables (all parameters are assumed time-independent), so that for example $N_H$ means $N_H(t)$. All parameters are summarized in Table 1. These equations can be expressed in an alternative form,

$$\dot{N}_H = r_{0,H} N_H \left(1 - \frac{N_H - c_{HL} N_L}{K_H}\right), \quad \text{eq. S2a}$$

$$\dot{N}_L = r_{0,L} N_L \left(1 - \frac{N_L - c_{LH} N_H}{K_L}\right), \quad \text{eq. S2b}$$

where $K_i = r_{0,i} / a_{ii}$ is the carrying capacity of species $i$ in the absence of the competitor, and $c_{ij} = a_{ij} / a_{ii}$ is the per-capita competitive effect of species $j$ on species $i$, scaled with respect to the within-species competitive effect of species $i$. In this formulation, the exponential growth is proportionally reduced by the (scaled) densities of the focal species and its competitor relative to the carrying capacity of the focal species.

Next, we consider effects of fishing and natural predation on crayfish dynamics. While several species like birds (e.g., herons) and fish (e.g., pikes) prey upon crayfish, we can assume that these predators do not specialize on crayfish; instead, as generalist predators they prey upon a more or less broad spectrum of species. As a consequence, such generalist predators will not show tightly coupled dynamics with their prey species, including crayfish, and the predators' dynamics do not have to be tracked in our community model (e.g. Turchin, 2003); this approach is in general also sensible for fishing. The usual way to account for the effects of generalist predators or fishing effects is to reduce the intrinsic growth rate of a focal species, e.g. an H-host, by a fixed effect, $r_H = r_{0,H} - f_H - p_H$. Here, $f_H = \rho_{HF} F$ is the effect on the growth rate of an H-host of $F$ fixed fishing units (e.g. density of anglers) with a per capita effect of $\rho_{HF}$. Analogously, $p_H = \rho_{HP} P$ is the effect on the growth rate of an H-host of $P$ fixed predator units (e.g. density of pikes) with a per capita effect of $\rho_{HP}$; $p_H$ could of course also be expressed as the sum of different predators feeding on a single H-host population. Note that the introduction of fishing and predation effects will also affect the carrying capacity, which now becomes $K_i = (r_{0,H} - f_H - p_H)/a_{ii}$.

The approach of fixing a predator's density to a constant will also, as an approximation, work for cases where a crayfish species preys for example upon a fish predator's eggs or juvenile stages. Crayfish are omnivorous (Longshaw & Stebbing, 2016), and thus it may be conceivable that this aspect could lead to "double predator-prey dynamics", where crayfish and fish are both prey and predator of each other. However, omnivorous species usually tend not to specialize on a single prey. Even if this were the case – with ensuing tightly coupled predator-prey dynamics – for our article's focus on epidemics what matters are long-term equilibrium densities of host and non-host species at the beginning of an epidemic (see section *S1. Eco-epidemiological dynamics: an overview*). Over the course of an epidemic's onset (on a time scale of days to weeks) changes in a crayfish host species density will have a negligible effect on a predator's density. This is because predator-prey dynamics of long-lived species will take time (on a time scale of months to years) to approach a new equilibrium state (Turchin, 2003). Thus, at an epidemic's onset fixing a predator's density to a constant value is justified as an approximation to more complex predator-prey dynamics.

Finally, to reflect the (potential) consequences of a pathogen on the whole host population, we will decrease the growth rate of an H-host by $\delta_H I_H$ because infected H-hosts will almost always die and therefore decrease total population size; as for an L-host, we do not introduce such decrease since L-hosts almost never die because of crayfish plague (see section *S3. Epidemiological dynamics*). In sum, population growth of an H-host and an L-host forming a community can be modelled as

$$\dot{N}_H = r_H N_H \left(1 - \frac{(N_H + c_{HL} N_L)}{K_H}\right) - \delta_H I_H, \qquad \text{eq. S3a}$$

$$\dot{N}_L = r_L N_L \left(1 - \frac{(N_L + c_{LH} N_H)}{K_L}\right). \qquad \text{eq. S3b}$$

To determine the disease-free equilibrium densities, we set $I_H = \dot{N}_H = \dot{N}_L = 0$ and solve for $N_H$ and $N_L$. Such competitive community can display different equilibrium states, and the general expressions for the species-specific equilibrium densities are

$$N_H^{eq} = \frac{K_H - c_{HL} K_L}{1 - c_{HL} c_{LH}}, \quad \text{eq. S4a}$$

$$N_L^{eq} = \frac{K_L - c_{LH} K_H}{1 - c_{HL} c_{LH}}. \quad \text{eq. S4b}$$

For coexistence to occur, it can be easily shown that the conditions $c_{HL} < K_H / K_L$ and $c_{LH} < K_L / K_H$ must hold (e.g. Kot, 2001). Recalling that both carrying capacities potentially depend on fishing and predation, for example the equilibrium density of the H-host ($N_H^{eq}$) will decrease if fishing or natural predation is present ($K_H$ decreases), and it will increase if for example fishing on the competitor (L-host) is intensified ($K_L$ decreases).

#### S3. Epidemiological dynamics

To construct the epidemiological sub-model (Fig. 3), we follow the well-established practice of subdividing a host population into several disease-related compartments (Keeling & Rohani, 2008). Usually, these compartments allow following the fate of susceptible individuals that get infected, without explicitly modelling the dynamics of the pathogen 'population'. Among the many variants of compartmental models, probably the most-often used one is the SIR-model, consisting of three compartments related to susceptible (S), infected (I), and recovered (R) individuals.

In general, and in the present case in particular because the ecological sub-model tracks the whole host population, the construction of the epidemiological sub-model should make sure that the sum of all host-specific compartments (their differential equations and the variables) result in the density and dynamics of the whole population (in our case the densities of H-hosts and L-hosts). For example, for an SIR-Model the whole population is the sum of all susceptible, infected, and recovered individuals, i.e. $N = S + I + R$.

In the present article, a main focus is on the condition for a crayfish plague outbreak to take off in a fully susceptible community. This focus allows simplifying the epidemiological sub-model by only considering epidemiologically relevant processes. One reason is that right at the beginning of an epidemic the host population (or community) is assumed to be at the disease-free equilibrium density: here, by definition the population growth rate (ecological sub-model) is zero. A second, related reason concerns the infected individuals: although the epidemic will alter the 'epidemiological composition' of the population (susceptible, infected, recovered individuals, or other compartments), right at the beginning of an outbreak the densities of infection-related compartments are (very) low, and thus their contribution to ecological processes are negligible.

*Multi-host compartmental model*

Our simplified epidemiological multi-host compartmental model, consisting of host- and compartment-specific differential equations (see below), is based on the following epidemiological mechanisms (see Table 1 for variables and parameters). With respect to living host individuals, we only track the dynamics of infected individuals ($I_i$) and of susceptible individuals ($S_i = N_i - I_i$) (cf. Bowers & Turner, 1997; Roberts & Heesterbeek, 2013), but the inclusion of additional compartments (e.g. for infected but not infectious individuals) would be straight-forward. The selection of the following mechanisms is based on previously published results (where available) or based on our own assessment:

- Susceptible individuals of an H-host species ($N_H - I_H$) are infected through the transmission of the pathogen (and disease) from the water, at a host-specific rate $\beta_H$ (Koivu-Jolma et al., 2023), and they remain infectious for a certain period ($\tau_H = \delta_H^{-1}$), after which they die. Because we are concerned with the beginning of an epidemic, the transmission modelled as a mass-action process is a sensible approximation. If the interest would be in an endemic equilibrium, then a more realistic function should be chosen (cf. Zhang et al., 2016).
- Susceptible individuals of an L-host species ($N_L - I_L$) are infected through the transmission of the pathogen (and disease) from the water, at a host-specific rate $\beta_L$ (Koivu-Jolma et al., 2023), and they remain infectious for a certain period ($\tau_L = \delta_L^{-1}$), after which they loose infectiousness. Because of our focus on the initial dynamics of an epidemic, we do not further specify their disease-related fate, e.g. life-long immunity, partial acquired immunity, or directly susceptible again. With regard to the functional form for the transmission process, see the preceding process.
- The bodies of deceased infected individuals of an H-host ($D_H$) will continue shedding pathogen spores into the water at a specific rate ($\sigma_D$) for a certain period ($\tau_D = \delta_D^{-1}$), after which they can be considered decomposed and non-infectious anymore (Makkonen et al., 2013).
- The exuviae of infected individuals of an L-host ($X_L$) will continue shedding pathogen spores into the water at a specific rate ($\sigma_X$) for a certain period ($\tau_X = \delta_X^{-1}$), after which they can be considered decomposed and non-infectious anymore (WOAH, 2016).
- The density of spores in the water ($W$) increases due to particle shedding of infected individuals of both host species, carcasses and exuviae, and decreases due to particle degradation (WOAH, 2016).

We did not include the following mechanisms:

- Compared to water-borne infection, the infection of susceptible individuals via contact with infected individuals – of the same or the second host species – has a negligible effect on overall pathogen transmission (Koivu-Jolma et al., 2023; WOAH, 2016).

- Compared to water-borne infection, the infection of susceptible individuals via scavenging-related contact with infected carcasses or exuviae has a negligible effect on overall pathogen transmission (Koivu-Jolma et al., 2023). Note that such infection events would not happen due to ingestion, but due to the spatial proximity during the scavenging event.
- Moulting of infected H-hosts is negligible (authors' assessment): an infected organism is likely to succumb to the disease without first moulting.
- Compared to the recovery rate of infected L-host individuals, infected individuals succumbing to the disease are rare (Oidtmann et al., 2006) and thus not included in the model.
- Uptake of fungal spores from the water during the transmission process is in principle expected to deplete the spore reservoir (cf. Cortez & Duffy, 2021). However, for crayfish – where the port of entry of the pathogen is via the exoskeleton and not via ingestion through active water filtration (see main text) – we can assume that a single spore upon being shed into the water is much more likely to be degraded or to leave the system (e.g. drifted away by water current) than to infect a susceptible crayfish. Mathematically, this translates to $-\left(\delta_W W + \beta_H \left(N_H - I_H\right) W\right) \approx -\delta_W W$ (eq. S5e).

The epidemiological dynamics are thus governed by

$$\dot{I}_H = \beta_H \left(N_H - I_H\right) W - \delta_H I_H, \qquad \text{eq. S5a}$$

$$\dot{I}_L = \beta_L \left(N_L - I_L\right) W - \delta_L I_L, \qquad \text{eq. S5b}$$

$$\dot{D}_H = \delta_H I_H - \delta_D D_H, \qquad \text{eq. S5c}$$

$$\dot{X}_L = \mu_L I_L - \delta_X X_L, \qquad \text{eq. S5d}$$

$$\dot{W} = \sigma_H I_H + \sigma_L I_L + \sigma_D D_H + \sigma_X X_L - \delta_W W. \qquad \text{eq. S5e}$$

*Basic reproduction numbers*

To derive the basic reproduction number of a community, $R_{0,C}$, the first step is to compute the Jacobian matrix, that is, differentiating all equations (eq. S5a-e) with respect to the epidemiological variables and evaluating it at the disease-free equilibrium densities, for which $I_H = I_L = D_H = X_L = W \approx 0$:

$$J = \begin{bmatrix} -\delta_H & 0 & 0 & 0 & \beta_H N_H^{eq} \\ 0 & -\delta_L & 0 & 0 & \beta_L N_L^{eq} \\ \delta_H & 0 & -\delta_D & 0 & 0 \\ 0 & \mu_L & 0 & -\delta_X & 0 \\ \sigma_H & \sigma_L & \sigma_D & \sigma_X & -\delta_W \end{bmatrix}. \qquad \text{eq. S6}$$

The Jacobian **J** can be expressed as the sum of a transmission matrix **B** and a transition matrix **T** (**J** = **B** + **T**), where

$$B=\begin{bmatrix} 0 & 0 & 0 & 0 & \beta_H N_H^{eq} \\ 0 & 0 & 0 & 0 & \beta_L N_L^{eq} \\ 0 & 0 & 0 & 0 & 0 \\ 0 & 0 & 0 & 0 & 0 \\ 0 & 0 & 0 & 0 & 0 \end{bmatrix} \qquad \text{eq. S7a}$$

$$T=\begin{bmatrix} -\delta_H & 0 & 0 & 0 & 0 \\ 0 & -\delta_L & 0 & 0 & 0 \\ \delta_H & 0 & -\delta_D & 0 & 0 \\ 0 & \mu_L & 0 & -\delta_X & 0 \\ \sigma_H & \sigma_L & \sigma_D & \sigma_X & -\delta_W \end{bmatrix}. \qquad \text{eq. S7b}$$

These two matrices are then used to compute the next-generation matrix with large domain as $\mathbf{M}_{L} = -\mathbf{B}\,\mathbf{T}^{-1}$ (Diekmann et al., 1990). This matrix has size 5 x 5, but because at the beginning of an epidemic infected individuals $I_i$ appear first, the relevant next-generation matrix (with small domain: see Diekmann et al., 1990, for further details) consists of the 2 x 2 entries of **M** in the top-left corner,

$$M=\begin{bmatrix} N_H^{eq}\dfrac{\beta_H(\delta_D\sigma_H+\delta_H\sigma_D)}{\delta_H\delta_D\delta_W} & N_H^{eq}\dfrac{\beta_H(\delta_X\sigma_L+\mu_L\sigma_X)}{\delta_L\delta_X\delta_W} \\ N_L^{eq}\dfrac{\beta_L(\delta_D\sigma_H+\delta_H\sigma_D)}{\delta_H\delta_D\delta_W} & N_L^{eq}\dfrac{\beta_L(\delta_X\sigma_L+\mu_L\sigma_X)}{\delta_L\delta_X\delta_W} \end{bmatrix}. \qquad \text{eq. S8}$$

The diagonal elements of **M** are the host species-specific $R_0$ values

$$R_{0,H}=N_H^{eq}\frac{\beta_H(\delta_D\sigma_H+\delta_H\sigma_D)}{\delta_H\delta_D\delta_W}, \qquad \text{eq. S9a}$$

$$R_{0,L}=N_L^{eq}\frac{\beta_L(\delta_X\sigma_L+\mu_L\sigma_X)}{\delta_L\delta_X\delta_W}, \qquad \text{eq. S9b}$$

while the off-diagonal elements contain terms summarizing epidemiological processes among host species. Note that, in isolation, each host would have its carrying capacity as equilibrium density instead of its equilibrium density adjusted for competition and predation (eq. S4a-b). Finally, the community-level basic reproduction number, $R_{0,C}$, corresponds to the leading eigenvalue (i.e. spectral radius) of **M**,

$$R_{0,C}=N_H^{eq}\frac{\beta_H(\delta_D\sigma_H+\delta_H\sigma_D)}{\delta_H\delta_D\delta_W}+N_L^{eq}\frac{\beta_L(\delta_X\sigma_L+\mu_L\sigma_X)}{\delta_L\delta_X\delta_W}=R_{0,H(C)}+R_{0,L(C)}. \quad \text{eq. S10}$$

Therefore, $R_{0,C}$ is the sum of the host species-specific $R_0$ values adjusted for competition and predation. This additive behaviour is caused by a lack of direct (i.e. contact) transmission among individuals of the two host species that share the same environment (Fig. 2a). These basic reproduction numbers – for an H-host, L-host, and a community – can be expressed in terms of the duration of infectiousness, so that $\tau_i=\delta_i^{-1}$. Then, as reported in Fig. 4 in the main text,

$$R_{0,H}=N_H^{eq}\beta_H(\sigma_H\tau_H+\sigma_D\tau_D)\tau_W, \quad \text{eq. S11a}$$

$$R_{0,L}=N_L^{eq}\beta_L(\sigma_L\tau_L+\mu_L\tau_L\sigma_X\tau_X)\tau_W \quad \text{eq. S11b}$$

#### S4. Basic reproduction numbers as a basis for management

*Sensitivity analysis*

A main utility of an $R_0$ expression is to quantitatively explore ways to keep or steer $R_0$ below the outbreak threshold $R_0$ = 1. Because in many cases parameter and variable values are not available – or at least are not precise enough to reflect the local specificities of a host population – an alternative way for $R_0$ to inform management decisions is by way of a sensitivity analysis (Caswell, 2019). Here, the main intent is to rank parameters and variables in terms of their relative effect on $R_0$: for example, if we change parameters and variables (one at the time) by *x*%, by how much would $R_0$ change? A simple approach to investigate this question is an elasticity analysis, and we start demonstrating it using the $R_0$ expression of an H-host (eq. S11a).

To derive elasticity expressions, first we differentiate $R_{0,H}$ with respect to parameters and variables ($\theta_j$) 'amenable' to management actions (see main text), leading to the sensitivity values $s_j=\partial R_{0,H}/\partial\theta_j$. The second step is to rescale these sensitivity values to give the elasticity values

$$e_{\theta_j}=\frac{\partial R_{0,H}}{\partial\theta_j}\frac{\theta_j}{R_{0,H}}. \quad \text{eq. S12}$$

An elasticity value tells us (to a first approximation) that slightly changing a parameter $\theta_j$, for example by 10%, will change $R_{0,H}$ by $100\cdot 0.1\cdot e_j$%. Thus, these values allow ranking parameters and variables – and therefore associated management actions – according to their 'efficiency' to change $R_{0,H}$. Because of the simple linear structure of $R_{0,H}$ or $R_{0,L}$ (eq. S11a-b), for example for $R_{0,H}$ in isolation we get the following elasticity values:

$$e_{N_H^{eq}} = e_{\tau_W} = 1, \quad \text{eq. S13a}$$

$$e_{\tau_H} = \frac{\sigma_H \tau_H}{\sigma_H \tau_H + \sigma_D \tau_D} < 1, \quad \text{eq. S13b}$$

$$e_{\tau_D} = \frac{\sigma_D \tau_D}{\sigma_H \tau_H + \sigma_D \tau_D} < 1. \quad \text{eq. S13c}$$

These results show that (i) reducing initial host density and reducing the persistence time of spores in the water body are equally efficient in reducing $R_{0,H}$ (proportionally speaking and disregarding the respective amount of resources needed), and (ii) reducing the infectious period of living and dead animals are both less efficient than the first two options. To more finely rank $e_{\tau_H}$ and $e_{\tau_D}$, the amount of total spores shed into the water by living and deceased animals would need to be estimated.

To use a sensitivity analysis for a crayfish community, as explained in the main text two situations should be considered separately. We start with an "emergency" situation, where mitigation actions would be applied not long before an outbreak is expected to happen, for example because of recently observed outbreaks in adjacent water bodies. Here, the time scale is expected not to be long enough for ecological processes like competition to react to the manipulation of host species densities as one potential action (see above). Thus, the sum of eq. S11a-b (see also Fig. 4) can be used by assuming that both host species densities are independent of each other (as opposed to eq. S4a-b). The parameters that potentially can be influenced by mitigation actions are thus $N_H^{eq}$, $N_L^{eq}$, $\tau_H$, $\tau_L$, $\tau_D$, $\tau_X$, $\tau_W$, with the aim of decreasing $R_{0,C}$ by decreasing equilibrium densities or durations of infectiousness. Following the calculation explained for a single host species above, the elasticity values are

$$e_{N_H^{eq}} = \frac{R_{0,H(C)}}{R_{0,C}} < 1, \quad \text{eq. S14a}$$

$$e_{N_L^{eq}} = \frac{R_{0,L(C)}}{R_{0,C}} < 1, \quad \text{eq. S14b}$$

$$e_{\tau_H} = \frac{N_H^{eq} \beta_H \sigma_H \tau_H \tau_W}{R_{0,C}} < 1, \quad \text{eq. S14c}$$

$$e_{\tau_L} = \frac{R_{0,L(C)}}{R_{0,C}} < 1, \quad \text{eq. S14d}$$

$$e_{\tau_D} = \frac{N_H^{eq} \beta_H \sigma_D \tau_D \tau_W}{R_{0,C}} < 1, \quad \text{eq. S14e}$$

$$e_{\tau_X} = \frac{N_L^{eq} \beta_L \mu_L \tau_L \sigma_X \tau_X \tau_W}{R_{0,C}} < 1, \quad \text{eq. S14f}$$

$$e_{\tau_W}=1. \qquad \text{eq. S14g}$$

These expressions show that reducing the persistence of spores in the water appears to be the most efficient option. Reducing host species densities in a community is now less efficient than the former option, compared to a single host population in isolation (see above). In addition, these elasticity expression allow ranking other options more finely: for example, for the L-host we have $e_{\tau_W}>e_{\tau_L}=e_{N_L^{eq}}>e_{\tau_X}$. A full ranking, nonetheless, will require a full parametrization.

As for the second situation with respect to a crayfish community to be managed, we assume that some management actions may be applied over a longer time period, without an expected incumbent outbreak. Because the manipulation of host densities could happen over a longer period of time, in contrast to an emergency situation (see above) competitive effects are now expected to react to the density manipulation. In addition to durations of infectiousness already analyzed in the previous example ($\tau_H$, $\tau_L$, $\tau_D$, $\tau_X$, $\tau_W$), we now add $K_L$ to this list which for example would be reduced in a sustained way by fishing.

The elasticity values for the duration of the infectious periods are the same as in eq. S14c-g. To compute the elasticity value for the carrying capacity of the L-host ($K_L$), we start again with the sum of eq. S11a-b as the community-level $R_0$ value (Fig. 4) and define $F_H:=\beta_H(\sigma_H\tau_H+\sigma_D\tau_D)\tau_W$ and $F_L:=\beta_L(\sigma_L\tau_L+\mu_L\tau_L\sigma_X\tau_X)\tau_W$, to get $R_{0,C}=N_H^{eq}F_H+N_L^{eq}F_L$. Next, we re-arrange the host equilibrium densities in the community (eq. S4a-b), resulting in $N_H^{eq}=K_HG^{-1}-c_{HL}K_LG^{-1}$ and $N_L^{eq}=K_LG^{-1}-c_{LH}K_HG^{-1}$, where we have defined $G:=1-c_{HL}c_{LH}$. Thus, community-$R_0$ can be rewritten as

$$R_{0,C}=\left(\frac{K_H}{G}-\frac{c_{HL}K_L}{G}\right)F_H+\left(\frac{K_L}{G}-\frac{c_{LH}K_H}{G}\right)F_L=K_H\left(\frac{F_H}{G}-\frac{c_{LH}F_L}{G}\right)+K_L\left(\frac{F_L}{G}-\frac{c_{HL}F_H}{G}\right). \qquad \text{eq. S15}$$

The elasticity value for the carrying capacity of the L-host (after simplification) reads

$$e_{K_L}=\left(1+\frac{K_H}{K_L}\frac{(F_H-c_{LH}F_L)}{(F_L-c_{HL}F_H)}\right)^{-1}. \qquad \text{eq. S16}$$

While a parametrization of eq. S16 is needed to robustly rank this management option (i.e. the sustained reduction of the invasive host species density) among all potential mitigation actions, from this expression an important caveat can be derived: density manipulation can backfire in terms of outbreak risk, despite the good intention of the management action. Because eq. S16 includes two differences (in brackets) involving the interaction of ecological and epidemiological processes, this elasticity value could in fact have negative sign: in such case, a desired *reduction* in $K_L$ would unintentionally lead to an *increase* of $R_{0,C}$. Whether or not this might be the case depends on the two ratios in eq. S16. The ratio of the two carrying capacities will have positive sign, and we can expect this ratio to be (much) smaller than 1. For example, the European noble crayfish in general attains a lower equilibrium density in isolation ($K_H$) compared to the respective $K_L$ value of several invasive crayfish species (Longshaw & Stebbing, 2016, and references therein). The second ratio in eq. S16 (containing the two differences), on the other hand, could have negative sign, in which case $e_{K_L}$ would become negative. Because $e_{K_L}$ becomes negative if the product of the two ratios is smaller than –1, a corollary can be derived: crayfish communities with host species attaining a similar carrying capacity in isolation ($K_i$) will *increase* the likelihood of increasing $R_{0,C}$ by *decreasing* $K_L$.

*Building $R_0$ ratios*

As we explain in the main text, an additional way of using $R_0$ values to inform management is to build $R_0$ ratios, with the aim of ranking single-host species or multi-host species communities in a region in terms of the likelihood and severity of an outbreak in case a pathogen invades. To illustrate the process of building such ratios, we choose a community consisting of an H-host and an L-host, and compare its $R_0$ value to the $R_0$ value of the same H-host alone, in the same environment. Using eq. S11a-b, the ratio reads

$$\frac{R_{0,C}}{R_{0,H}}=\frac{R_{0,H(C)}+R_{0,L(C)}}{R_{0,H}}=\frac{N_H^{eq}}{K_H}+\frac{N_L^{eq}}{K_H}\frac{\beta_L}{\beta_H}\frac{\sigma_L\tau_L+\mu_L\tau_L\sigma_X\tau_X}{\sigma_H\tau_H+\sigma_D\tau_D}, \qquad \text{eq. S17}$$

where for this illustrative example $\tau_W$ cancels out. This expression can be rearranged, with the aim of having all parameters and variables as ratios, too. These ratios will then allow gauging the values of parameters and variables in a relative way. The rearranged form of eq. S17 (see Box 1) reads

$$\frac{R_{0,C}}{R_{0,H}}=\frac{N_H^{eq}}{K_H}+\frac{N_L^{eq}}{K_H}\frac{\beta_L}{\beta_H}\frac{\sigma_X\tau_X}{\sigma_D\tau_D}\frac{\left(\frac{\tau_L}{\tau_\mu}+\frac{\sigma_L\tau_L}{\sigma_X\tau_X}\right)}{\left(1+\frac{\sigma_H\tau_H}{\sigma_D\tau_D}\right)}. \qquad \text{eq. S18}$$

Bozzuto C, Schmidt-Posthaus H, Adrian-Kalchhauser I, Pisano SRR (2024): Towards an eco-epidemiological framework for managing freshwater crayfish communities confronted with crayfish plague. Preprint *bioRxiv*.

#### References Supporting Information

Bowers, R. G., & Turner, J. (1997). Community Structure and the Interplay between Interspecific Infection and Competition. *Journal of Theoretical Biology*, *187*(1), 95–109. https://doi.org/10.1006/jtbi.1997.0418

Caswell, H. (2019). *Sensitivity Analysis: Matrix Methods in Demography and Ecology*. Springer Nature. https://doi.org/10.1007/978-3-030-10534-1

Cortez, M. H., & Duffy, M. A. (2021). The Context-Dependent Effects of Host Competence, Competition, and Pathogen Transmission Mode on Disease Prevalence. *The American Naturalist*, 198(2), 179–194. https://doi.org/10.1086/715110

Diekmann, O., Heesterbeek, J. A. P., & Metz, J. A. J. (1990). On the definition and the computation of the basic reproduction ratio R0 in models for infectious diseases in heterogeneous populations. *Journal of Mathematical Biology*, *28*(4), Article 4. https://doi.org/10.1007/BF00178324

Keeling, M. J., & Rohani, P. (2008). *Modeling Infectious Diseases in Humans and Animals*. Princeton University Press.

Koivu-Jolma, M., Kortet, R., Vainikka, A., & Kaitala, V. (2023). Crayfish population size under different routes of pathogen transmission. *Ecology and Evolution*, *13*(1), Article 1. https://doi.org/10.1002/ece3.9647

Kot, M. (2001). *Elements of Mathematical Ecology*. Cambridge University Press. https://doi.org/10.1017/CBO9780511608520

Longshaw, M., & Stebbing, P. (2016). *Biology and Ecology of Crayfish*. CRC Press.

Makkonen, J., Strand, D. A., Kokko, H., Vrålstad, T., & Jussila, J. (2013). Timing and quantifying Aphanomyces astaci sporulation from the noble crayfish suffering from the crayfish plague. *Veterinary Microbiology*, *162*(2), 750–755. https://doi.org/10.1016/j.vetmic.2012.09.027

Oidtmann, B., Geiger, S., Steinbauer, P., Culas, A., & Hoffmann, R. W. (2006). Detection of Aphanomyces astaci in North American crayfish by polymerase chain reaction. *Diseases of Aquatic Organisms*, *72*(1), 53–64. https://doi.org/10.3354/dao072053

Roberts, M. G., & Heesterbeek, J. A. P. (2013). Characterizing the next-generation matrix and basic reproduction number in ecological epidemiology. *Journal of Mathematical Biology*, *66*(4), Article 4. https://doi.org/10.1007/s00285-012-0602-1

Roberts, M. G., & Heesterbeek, J. a. P. (2018). Quantifying the dilution effect for models in ecological epidemiology. *Journal of The Royal Society Interface*, *15*(140), Article 140. https://doi.org/10.1098/rsif.2017.0791

Roberts, M. G., & Heesterbeek, J. a. P. (2020). Characterizing reservoirs of infection and the maintenance of pathogens in ecosystems. *Journal of The Royal Society Interface*, *17*(162), Article 162. https://doi.org/10.1098/rsif.2019.0540

Roberts, M. G., & Heesterbeek, J. a. P. (2021). Infection dynamics in ecosystems: On the interaction between red and grey squirrels, pox virus, pine martens and trees. *Journal of The Royal Society Interface*, *18*(183), Article 183. https://doi.org/10.1098/rsif.2021.0551

Turchin, P. (2003). *Complex Population Dynamics: A Theoretical/empirical Synthesis*. Princeton University Press.

WOAH. (2016). *Manual of diagnostic tests for aquatic animals* (Seventh edition). https://portals.iucn.org/library/node/48451

Zhang, L., Wang, Z.-C., & Zhang, Y. (2016). Dynamics of a reaction–diffusion waterborne pathogen model with direct and indirect transmission. *Computers & Mathematics with Applications*, *72*(1), 202–215. https://doi.org/10.1016/j.camwa.2016.04.046
